# Thermal and Nutrient Stress Drove Permian-Triassic Marine Extinctions

**DOI:** 10.1101/2023.09.08.556797

**Authors:** William J. Foster, Anja B. Frank, Qijan Li, Silvia Danise, Xia Wang, Jörn Peckmann

## Abstract

The Permian-Triassic mass extinction coincides with extensive environmental changes (i.e., thermal stress, deoxygenation and potentially ocean acidification), but the primary drivers of extinction in them marine realm are currently unknown. To understand which factors caused extinctions, we quantitatively investigated the relationship between geochemical proxies and fossil record at the most intensively-studied locality for this event, the Meishan section, China. We found that δ^18^O_apatite_ (paleotemperature proxy) and δ^114^Cd (primary productivity proxy) best explain changes in species diversity and composition at Meishan’s paleoequatorial setting. These findings suggest that the physiological stresses induced by ocean warming and nutrient availability played a predominant role in driving equatorial marine extinctions during the Permian-Triassic event. This research enhances our understanding of the interplay between environmental changes and extinction dynamics during a past climate crisis.

**One-Sentence Summary:** Ocean warming and nutrient availability were key drivers of equatorial marine extinctions during the Permian-Triassic mass extinction.

## Main Text

The Permian-Triassic climate crisis is reflected by an exceptionally rapid warming (around 8-10°C in 10-20 ka at low latitudes) in the latest Permian into the Early Triassic (late Griesbachian) (*1*). This climate crisis is thought to have been caused by the simultaneous eruptions of the Siberian Traps Large Igneous Province and combustion of organic-rich 5 sedimentary rocks (*2*). This climate crisis is also associated with the Permian-Triassic mass extinction, the most catastrophic mass extinction on Earth, which was highly selective against taxonomic groups that dominated pre-extinction marine communities, with an estimated loss of 96% of species (*3*). A number of environmental changes have been proposed as being the main drivers of marine extinction during the Permian-Triassic transition (most notably deoxygenation, 10 thermal stress, and ocean acidification (*3-4*)), yet it is not known exactly which factors played a major role in causing the biodiversity crisis. In addition, because during the event, multiple environmental perturbations occur simultaneously, it is difficult to disentangle which specific environmental changes were most significant in causing the extinctions. This lack of understanding is also due to the poor geographical coverage of continuous Permian-Triassic 15 successions and the small number of sections that have been investigated at a high-resolution with multiple proxies for environmental and biodiversity changes.

Along the Meishan Hill, Zhejiang, China, a Permian-Triassic succession expands 2 km laterally and has been the subject of many paleontological and geochemical studies (5), which combined make the Meishan sections the only place that can currently be quantitatively investigated to 20 better understand which environmental proxies relate to biodiversity loss. In addition, Meishan’s Permian-Triassic succession has a clear stratigraphic scheme with each bed and subbed numbered allowing accurate correlations between studies performed over the last 3 decades. During the Permian-Triassic transition, the Meishan sections represent an outer slope setting in an equatorial (ca. 20°N) epicontinental sea (*6*). This means that the Meishan sections can provide 25 an analog into the causes of extinction during an extreme climate crisis for equatorial, shallow marine ecosystems (i.e., for the worst-case RCP scenario which predict a total temperature increase of more than 4.5°C by 2100 (*7*)). Here, we have created a database of the fossil record (603 species from 6457 occurrences) and 18 geochemical proxies (Table S1) for environmental change from the Meishan sections to 1) define the timing of extinction among different marine 30 taxa, and 2) quantitatively investigate which environmental changes associated with the climate crisis best explain the marine extinctions.

## Results

### Nature of the mass extinction event

The nature of the Permian-Triassic mass extinction is hotly debated and has been interpreted 35 either as a single pulse/ interval or a multiple-pulsed extinction (*8-9*). Quantifying the number of pulses of extinction, which takes into account confidence intervals of stratigraphic ranges (see supplemental material), demonstrates that the nature of the mass extinction is complex and varies between different phyla. Near the Permian/Triassic boundary, the traditional extinction horizon (bed 25) (*9*) marks a large number of last occurrences (LAD), leading to composition shifts 40 (Figs. 1, S1-S5), however our analysis shows that a single pulse of extinction (the final LADs) occurs at bed 28 for mollusks, and bed 29a for foraminifera, brachiopods, and conodonts (Fig. 1). Ostracods, instead, record two earlier pulses of extinction at beds 23a and 24d (Fig. 1), the latter coinciding with a sequence boundary. The species richness of the remaining groups are too low to detect the timing of extinction, but the highest occurrences do not occur above bed 27c. This 45 suggests that the nature of the mass extinction event at Meishan, except for ostracods, is best characterized as an extinction interval (51 cm), from beds 25 to 29a (*C. meishanensis – I. isarcia* conodont zones). Such interpretation is also relatively consistent with a stark reduction in bioturbation and tiering depth at the base of bed 25 (*10*). Whereas ostracods record an earlier major extinction interval from beds 22-23a and a subsequent minor pulse at bed 24d (*11*), suggesting that this group of organisms were more sensitive to initial environmental changes or 5 responded to different environmental changes prior to the main extinction interval. When the timing of extinction is investigated including all of the species, the mass extinction event is consistent between the different data splits, with a 2-pulse event at beds 23a and 29a, with bed 23a reflecting the selective extinction of ostracods (Table S2). In addition to the extinction interval that spans the Permian/Triassic boundary, ostracods, mollusks, brachiopods, and 10 conodonts record a minor extinction pulse earlier in the Changhsingian too (Fig. 1).

**Fig. 1.**
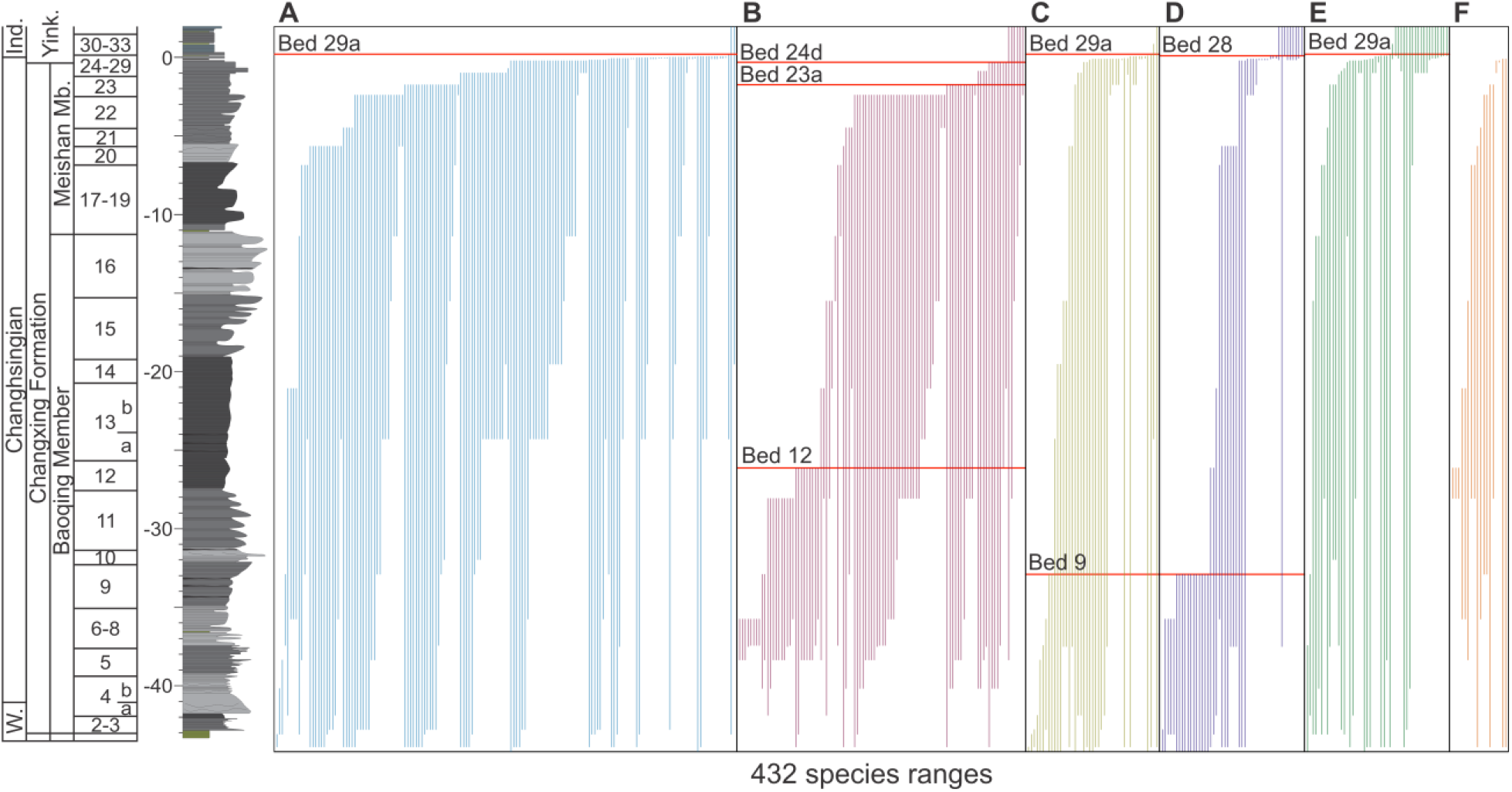
Stratigraphic ranges of fossil species (vertical lines) from the Meishan sections. Stratigraphic ranges of (**A**) foraminifera, (**B**) arthropods (all ostracods, except 1 trilobite species), (**C**) brachiopods, (**D**) mollusks, **(E**) conodonts, and (**F**) other (includes: bryozoans, corals, 15 calcareous algae, and *Tubiphytes*). Quantitatively determined extinction pulses for each phylum indicated (horizontal red line). Singletons are excluded from the figure and from determining the number of extinction pulses. Bed numbers and sedimentology follows Zhang et al. (*12*) follow Yin et al. (*13*). 0 meters is taken as the base of bed 27c, which is the biostratigraphic position of the Permian/Triassic boundary that is defined by the first appearance of *Hindeodus parvus. W. =* 20 *Wuchiapingian, Ind. = Induan, Yink. = Yinkeng Formation*.

These changes are also reflected by the breakpoints and rapid decline in species diversity at the Meishan sections at beds 22, 24e, and 29a (Fig. 2). Lithology changes and sequence stratigraphic boundaries play a role in determining the stratigraphic position of last occurrences (*14*). At Meishan, the extinction interval includes a lithostratigraphic boundary at the base of bed 25, and 25 a transgressive surface at bed 27a, suggesting that sedimentological changes affect our interpretations of the nature and timing of the extinction. Despite that, radiometric dating suggests that bed 25 to 28 only represents 60 ±48 ka (*15*) and any hiatuses associated with sequence stratigraphic surfaces during the extinction interval are of a relatively short duration.

**Fig. 2.**
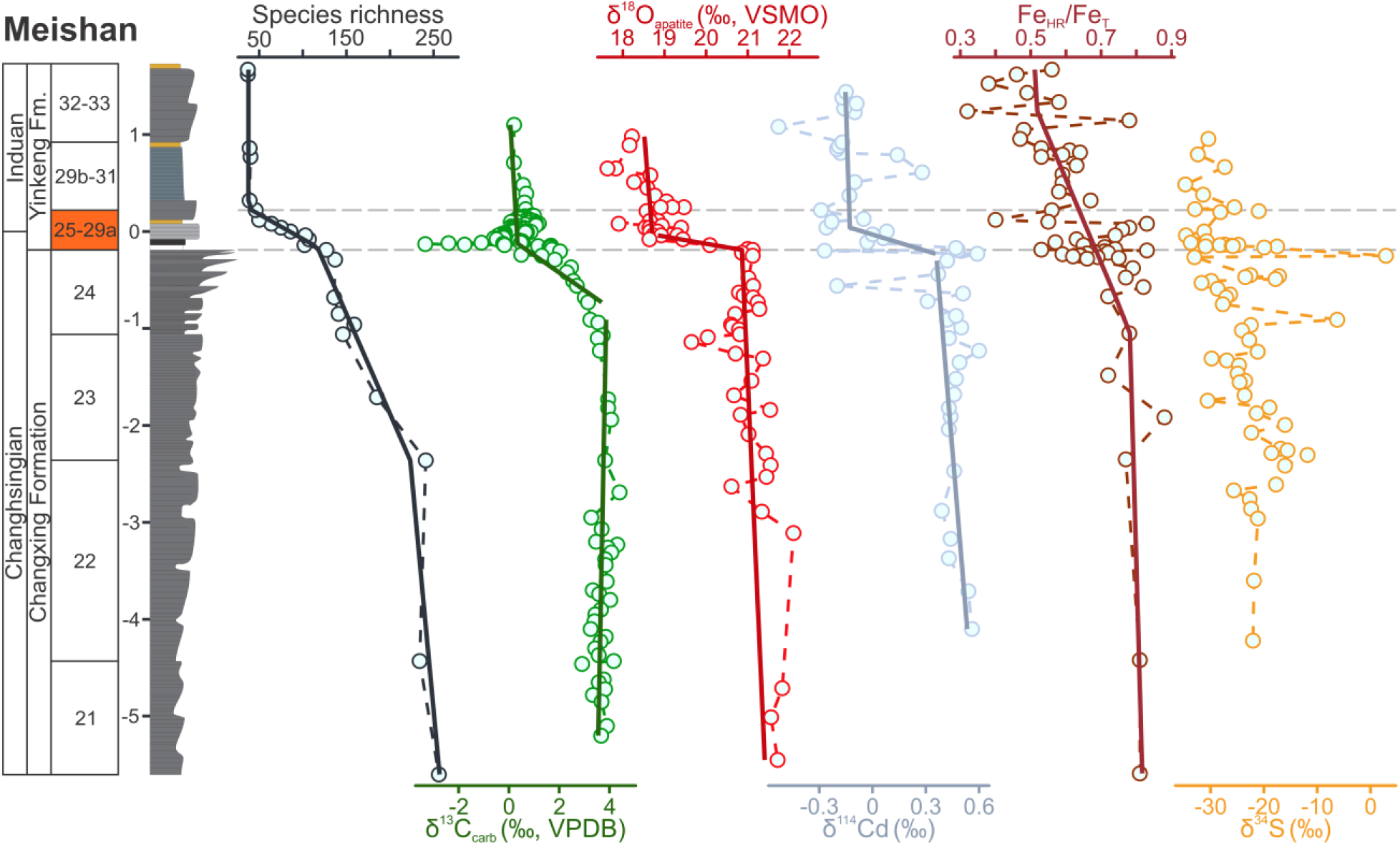
Stratigraphic correlation of selected paleoenvironmental proxies with species 10 diversity at the Meishan sections, South China, with segmented regression lines overlain. δ^13^C_carb_ from Shen et al. (*17*), δ^18^O_apatite_ (VSMO) from Chen et al. (*18*), δ^114^Cd isotopes from Zhang et al. (*16*), Fe_HR_/Fe_T_ from Xiang et al. (*19*), δ^34^S from Shen et al. (*20*). The main extinction interval (beds 25-29a) is highlighted in orange and with two horizontal dashed lines. Note: only paleoenvironmental proxies that showed significant relationships with diversity are included, for 15 a full figure with all the paleoenvironmental proxies see Figs. S6-S8.

Changes in species diversity, in particular the earlier onset of ostracod extinctions, are problematic when trying to compare extinctions with geochemical proxies. This is because many of the proxies that have been investigated at the Meishan sections only span a short interval, e.g. δ^114^Cd only spans beds 22-33 (*16*), after species diversity has already started to decline (Fig. 2). 5 Analyses linking geochemical and fossil data were, therefore, restricted to beds 22-29a. Investigations did not extend beyond bed 29a, because of protracted low diversity after the extinction interval due to delayed recovery rather than to environmental conditions.

### Quantifying the causes of extinction

Quantifying the causes of extinction is complex as environmental changes will manifest with different patterns and may be reflected as either state-shifts, anomalies, or trends that can be 20 associated with biodiversity dynamics. For example, the investigated proxies associated with volcanism, e.g., Hg/TOC, δ^66^Zn and δ^187^Os, are expected to appear as anomalies or spikes. Hg/TOC, δ^66^Zn and δ^187^Os show anomalies that coincide with the onset of the mass extinction interval (Fig. S8), with the Hg/TOC, δ^66^Zn, and δ^187^Os anomalies from bed 24b-24e being interpreted to reflect volcanism associated with the Siberian Traps coming along with input of volcanic ashes (*21-23*).

A segmented regression analysis, which can be used to quantify significant temporal shifts in proxies (i.e., state-shifts), recognizes significant changes for δ^13^C_carb_, δ^18^O_apatite_, δ^114^Cd, and δ^15^N 5 at the onset of the extinction interval (Fig. 2, Figs S6-8). In addition, TOC shows a breakpoint at bed 22 (Fig. S7), corresponding with the main extinction pulse of ostracods (Fig. 1). Whereas Th/U_apatite_ ratios show a state shift at bed 29 (Fig. S7), with an interpreted state shift from oxic to anoxic conditions (*24*), and corresponding with a plateau of low-diversity.

One issue with comparing the different proxies and changes in species richness or incidence data 10 is the difference in the resolution of the different datasets. To allow for statistical exploration of the data, the data was aggregated to the same resolution of the species dataset, i.e., bed and subbed-level resolution. In addition, not all of the beds record proxy values and, therefore, data were extrapolated using the segmented regression curves for each proxy (Figs. S9-S10). Another issue is that many of the different geochemical proxies correlate with one another, making it 15 difficult to disentangle if the proxy is robust enough to interpret environmental changes, if an environmental change is causing a decline in diversity, or if both diversity and proxy dynamics have a common cause. A correlation plot shows that δ^13^C_carb_, δ^18^O_apatite_, δ^114^Cd, and δ^15^N are significantly correlated (Figs. S11-13), which suggests these proxies share a common cause.

Many of the proxies show a correlation with changes in species richness (Tab. S3). A Poisson 20 regression model was performed to identify which proxies best explain the diversity dynamics. Value-inflated factors show that the correlation between δ^13^C_carb_, δ^18^O_apatite_, δ^114^Cd, and δ^15^N significantly affects the quality of the model. For this reason, and because the δ^15^N data is at a low resolution, δ^15^N was dropped from the model, whereas for δ^114^Cd and δ^18^O_apatite_ two separate models were run. The generalized linear models show that δ^13^C_carb_, δ^18^O_apatite_, δ^114^Cd, and 25 δ^44^Ca_apatite_ have significant relationships with changes in species diversity at Meishan (Tab. 1). In addition, and no proxies showed a significant relationship between proxy variance and extinction rate.

**Table 1.** Generalized linear model of significant environmental variables (geochemical proxies) and changes in diversity. Model selection was based on proxies that showed consistent and significant linear relationships with diversity (Tab. S3). Note: δ^114^Cd and δ^15^N were dropped from the first model because they showed a significant correlation with δ^18^O that negatively 5 impacted the model (Supp. mat). A second model swapping δ^18^O and δ^114^Cd was done to investigate the best model.

| Model | Parameter | Estimate | 95% Confidence intervals |  | t-value | p-value |
| --- | --- | --- | --- | --- | --- | --- |
|  |  |  | 2.5% | 97.5% |  |  |
| Species<br>Diversity<br>pseudo-$R^2 = 0.62$ | (Intercept) | 0.40 | -1.35 | 2.15 | 0.44 | 0.658 |
| | $\delta^{13}\text{C}_{\text{carb}}$ | 0.06 | 0.01 | 0.11 | 2.49 | **0.001** |
| | $\delta^{18}\text{O}_{\text{apatite}}$ | 0.25 | 0.17 | 0.32 | 6.38 | **< 0.001** |
| | $\delta^{44}\text{Ca}_{\text{apatite}}$ | 0.94 | 0.22 | 1.67 | 2.55 | 0.011 |
| Species<br>Diversity<br>pseudo-$R^2 = 0.59$ | (Intercept) | 5.71 | 5.13 | 6.30 | 19.18 | **< 0.001** |
| | $\delta^{13}\text{C}_{\text{carb}}$ | 0.03 | -0.03 | 0.09 | 1.08 | 0.279 |
| | $\delta^{114}\text{Cd}$ | 0.92 | 0.57 | 1.28 | 5.09 | **< 0.001** |
| | $\delta^{44}\text{Ca}_{\text{apatite}}$ | 1.61 | 0.92 | 2.31 | 4.55 | **< 0.001** |

A partial distance-based redundancy analysis (partial-dRDA) was undertaken to investigate the changes in fossil incidence data for beds 22 to 29a and changes in geochemical proxies (Fig. 3). Once more, value-inflated factors show that the correlation between δ^13^C_carb_, δ^18^O_apatite_, δ^114^Cd, 10 and δ^15^N significantly affects the quality of the model. δ^114^Cd and δ^15^N were, therefore, dropped from the model. The partial-dRDA showed that δ^13^C_carb_, δ^18^O_apatite_, and ^87^Sr/^86^Sr best explained changes in the incidence data. Swapping δ^114^Cd with δ^18^O_apatite_ shows that δ^18^O_apatite_ is a more significant proxy in explaining incidence data dynamics. When only the significant factors are included in the partial-dRDA model, only δ^13^C_carb_ and δ^18^O_apatite_ record significant relationships 15 (Fig. 3). It is also evident that the fossil incidence data cluster according to lithology (Fig. 3), highlighting how lithological changes reflect changes in the environment affecting species loss.

**Fig. 3.**
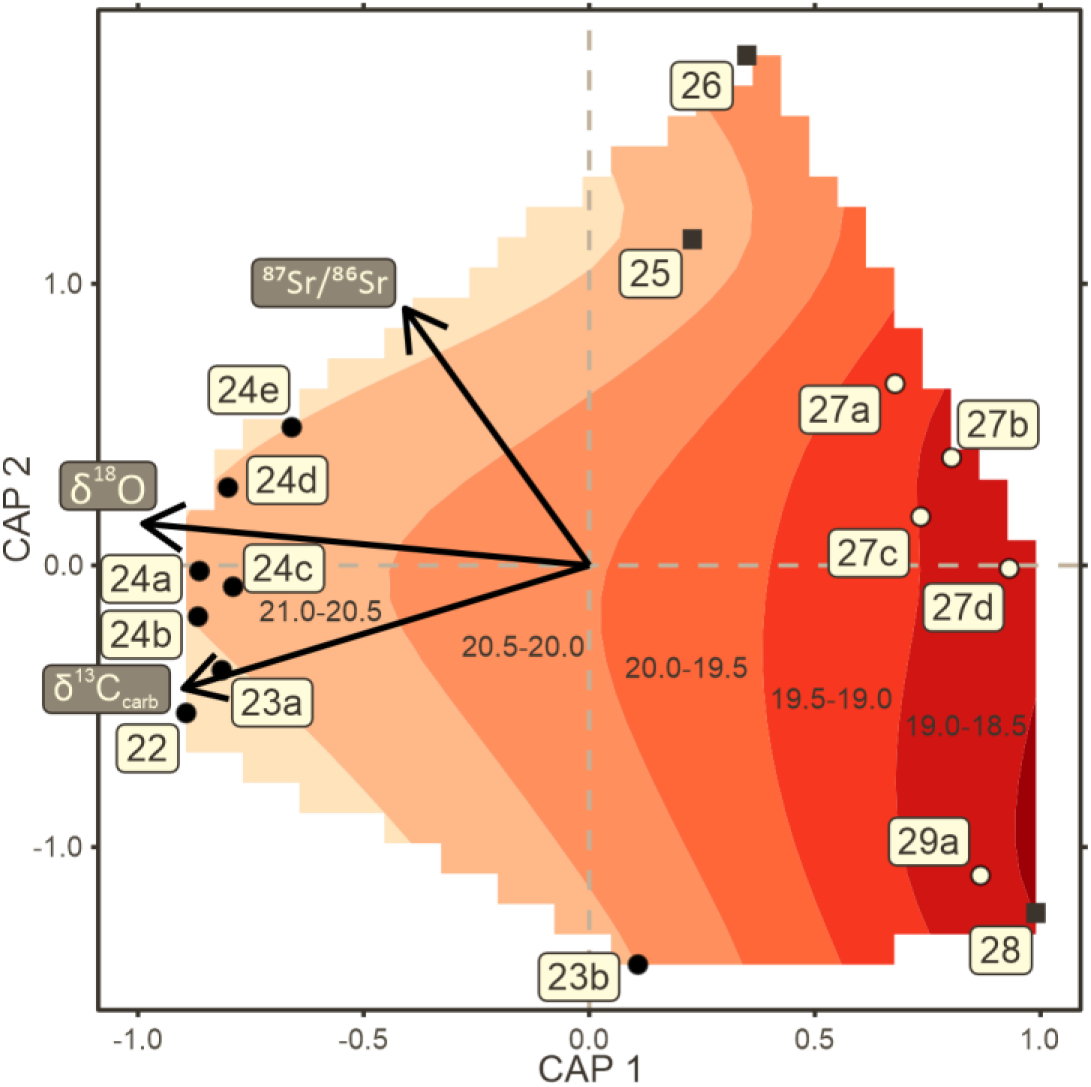
Partial Distance-based Redundancy Analysis (capscale) for fossil assemblages and geochemical proxies from the Meishan sections. Included vectors are the geochemical proxies that were determined as having a significant relationship with the fossil assemblages. Sample point shapes relate to bed lithology: filled circles = limestone, open circles = silty limestone, and filled squares = clay. The bed numbers for each assemblage are indicated and only beds 22-29a are included due to limited coverage of geochemical proxies at the Meishan sections. Smooth contour of the oxygen isotope values underlay the ordination plot to demonstrate the relationship with the fossil assemblages.

## Discussion

Due to the large suite of geochemical proxies investigated for the Meishan sections, a high number of different environmental changes have been proposed as possible causes of the Permian-Triassic mass extinction. Our quantitative analysis combining diversity and proxy data, demonstrates that δ^13^C_carb_, δ^18^O_apatite_, and δ^114^Cd are key for understanding the cause(s) of the mass extinction event. The geochemical signature of these proxies is mostly generated in the euphotic zone with a high chance for transfer into sediments without much alteration, suggesting that the relationships between these proxies and the fossil record reflect past environmental-life interactions. In addition, the lack of relationship between the fossil incidence data and the geochemical proxies that are affected by lithology changes suggests that our interpretations are robust. δ^13^C_carb_ is often interpreted as reflecting the release of large quantities of isotopically light carbon into the atmosphere, changes in primary productivity, and changes in carbon burial rates (*25*). δ^13^C_carb_ can, therefore, signify an environmental disturbance and even the trigger of the mass extinction (*26*), but cannot be inferred to identify the underlying environmental changes that drove species to extinction. In addition, δ^13^C_carb_, δ^18^O_apatite_, and δ^114^Cd are significantly correlated (Fig. S11), which we infer as being impacted by a common cause.

A negative excursion in δ^18^O_apatite_ is interpreted to reflect a rapid, 8-10°C, warming associated with the mass extinction event at Meishan (*1*) and is consistently the best explanatory factor for diversity dynamics at Meishan (Tab. 1, Fig. 3). Thermal stress is understood to limit the performance of aerobic marine organisms, because the pejus temperature is close to the temperature optimum on the upper thermal limit, and increasing temperatures beyond a marine organisms optimum temperature range rapidly leads to a reduction in the aerobic scope of marine organisms (excess capacity supporting activity, growth and reproduction) (*27*). In addition, the paleoequatorial setting of the Meishan section also suggests that this locality would also have experienced some of the highest climate velocities (*sensu 28*) with Earth system models for the Permian-Triassic mass extinction, with the consequent loss of well-oxygenated habitats for marine ecosystems (*29*). The significant relationship between δ^18^O_apatite_ with diversity and compositional changes recorded here supports the interpretation that temperature driven hypoxia was fundamental in causing equatorial extinctions/extirpations during the Permian-Triassic mass extinction.

δ^18^O_apatite_ is also significantly correlated with δ^114^Cd and δ^15^N, where both δ^114^Cd and δ^15^N record negative excursions that are interpreted to reflect primary productivity dynamics (*16, 30*). δ^114^Cd reflects nutrient utilization by phytoplankton and is an indirect proxy for primary productivity. At Meishan a negative excursion in δ^114^Cd coincides with the mass extinction interval reflecting a collapse in primary productivity (Fig. 2; *16*), which is also reflected in the major extinction of radiolarians (the primary fossil record of plankton biodiversity) (*31*). A stark reduction in primary productivity would be catastrophic for marine ecosystems because the cascading effect of extinction causes knock-on effects on species populations in successive layers of the marine food web (*32*). In addition, the aerobic metabolism is not only affected by thermal stress, but also nutritional stress, which can exacerbate the effects of climate change in marine ectotherms (*33*). The impacts on the aerobic metabolism of marine organisms can also be inferred from the observed decrease in the body-size of surviving taxa (*34*). In the equatorial paleosetting of the Meishan sections, both thermal stress and a collapse in primary productivity are interpreted to best explain the extinctions.

It has been shown that despite inter-specific differences, there are clear differences in hypoxia tolerance amongst higher taxa (*35*). Ostracods have the least tolerance to hypoxia, and their earlier onset of extinction at the Meishan section also corresponds to the initial changes in the δ^18^O_apatite_ negative excursion (Fig. 2). δ^18^O_apatite_ dynamics have been divided into two phases associated with different rates and magnitude of warming (*36*), with the first phase, coinciding with the major extinction of ostracods, being slower and of smaller magnitude. Pre-extinction changes in δ^18^O_apatite_ in equatorial settings have also been related to body size changes in ammonoids (*37*) and also correspond to pre-extinction changes in brachiopod assemblages (*38*). This suggests that, pre-extinction slower warming and the following rapid warming, led to the different timing of extinction between different marine organisms, depending on their sensitivity to temperature and oxygen-concentration changes. Although the Meishan sections have been the subject of several paleoredox studies, utilizing various proxies resulting in different redox interpretations (*19-21, 30*), the timing and extent of anoxic episodes are too poorly constrained (supplemental material) to adequately assess the role anoxia played in the Permian-Triassic mass extinction.

Despite the intense geochemical and paleontological research on the Meishan sections, this study also highlights some shortcomings that must be addressed in future research. The restriction of investigations of geochemical proxies over a short-interval at the Permian/Triassic boundary hinders our ability to understand how environmental conditions evolved over the Changhsingian and how that relates to the climate crisis (e.g., δ^44^Ca; *39*). Therefore, even though δ^44^Ca, a potential proxy for ocean pH, is recorded as having a significant relationship with changes in species richness (Tab. 1), the short record still makes this interpretation equivocal. In addition, the lack of abundance data and other ecological data from paleontological studies means it is not yet possible to investigate the ecological impacts of the Permian-Triassic climate crisis beyond the timing of extinction (e.g., *11, 40*). Therefore, a number of ecological changes, such as changes in relative abundance, dominance, or body size and how they relate to environmental changes cannot be explored. Finally, the cause(s) of extinction are expected to vary on various spatial scales, and therefore more high-resolution studies from other sections and regions are also required. Despite these short-comings, our statistical analysis demonstrates that the extreme impact of environmental changes on the aerobic metabolism of marine ectotherms and cascading effects of extinction best explain the cause of extinction in an epicontinental, equatorial settings.

## Acknowledgments

We would like to thank Steve Wang (Swathmore College) for his work developing the parallelization of the extinction pulse algorithm. We would like to thank all the scientists who have entered fossil occurrence data into both the Geobiodiversity database and Paleobiology database. We would like to thank the CEN-IT at Universität Hamburg for access to the computer cluster required for our analyses.

## Author contributions

Conceptualization: WJF

Methodology: WJF, ABF, QL, SD, SW

Investigation: WJF, ABF, SD, XW

Visualization: WJF

Funding acquisition: WJF

Writing – original draft: WJF

Writing – review & editing: all authors

